## Supplementary Materials for "Molecular dynamics simulation analysis of structural dynamic cross correlation induced by odorant hydrogen-bonding in mouse eugenol olfactory receptor^*^"

#### Supplementary Figure S1

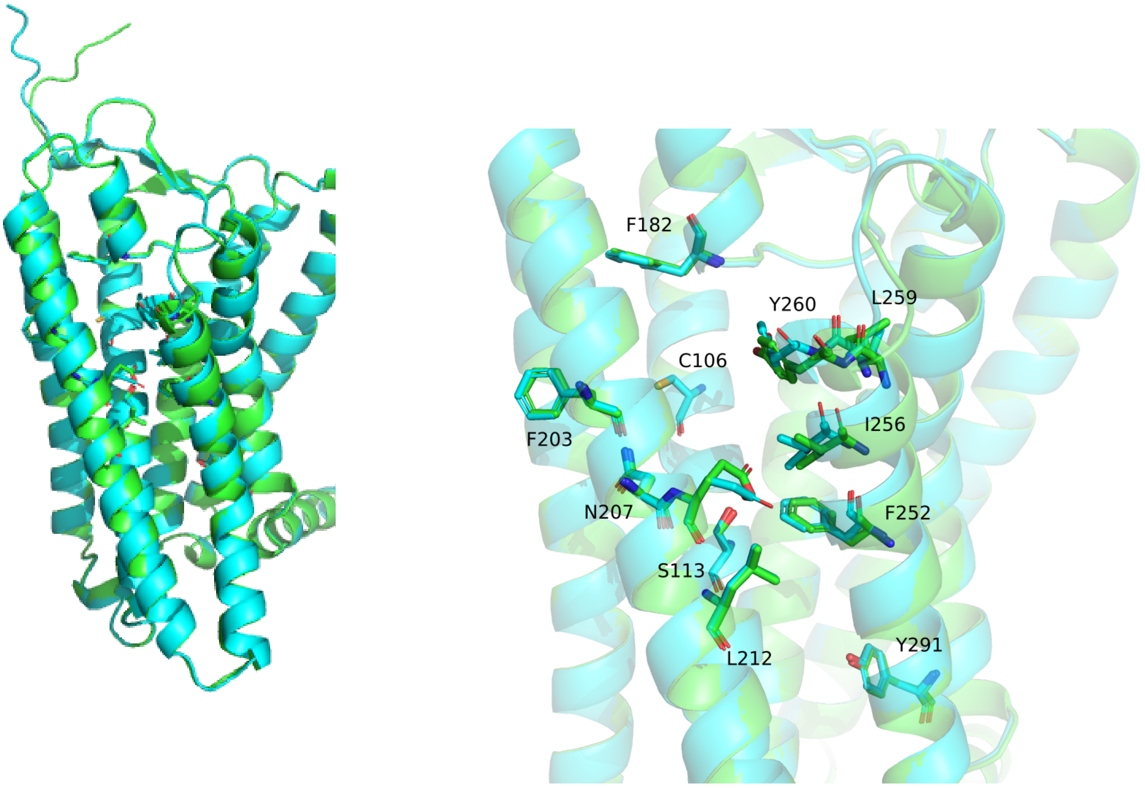


**Supplementary Figure S1** Comparison between the present result from the ColabFold server (green, accessed on July 8, 2022) and the UniProt entry Q920P2 (cyan, AF-Q920P2-F1-model_v4.pdb). The right panel corresponds to Figure 1 in the main text.

### Supplementary Figure S2

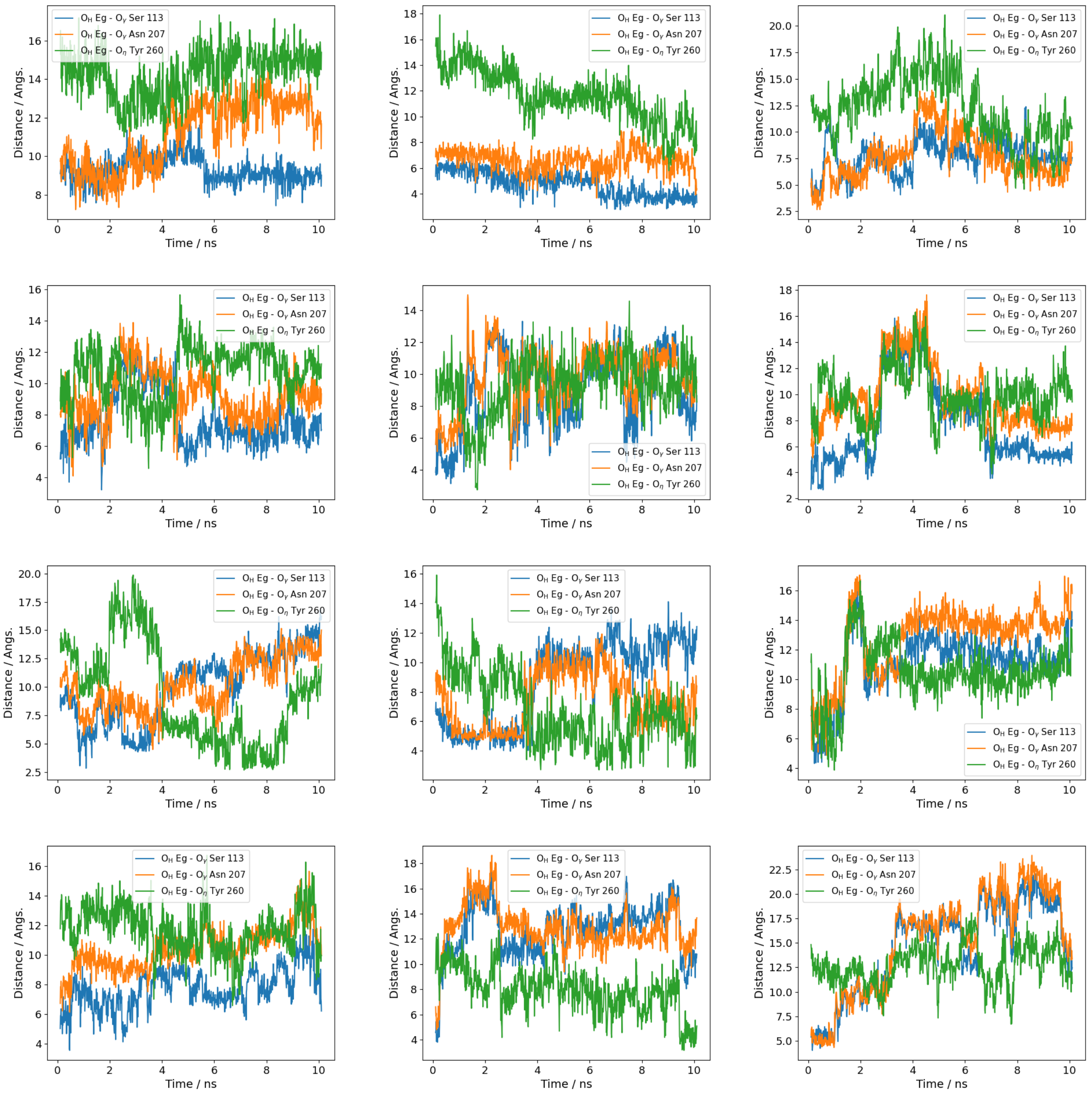


**Supplementary Figure S2** Eighteen trajectories (in addition to the two in Figure 3 in the main text) of the distance between $O_H$ atom of eugenol and O atoms of Ser113, Asn207, and Tyr260 starting from the docking configurations. (Continue to next page.)

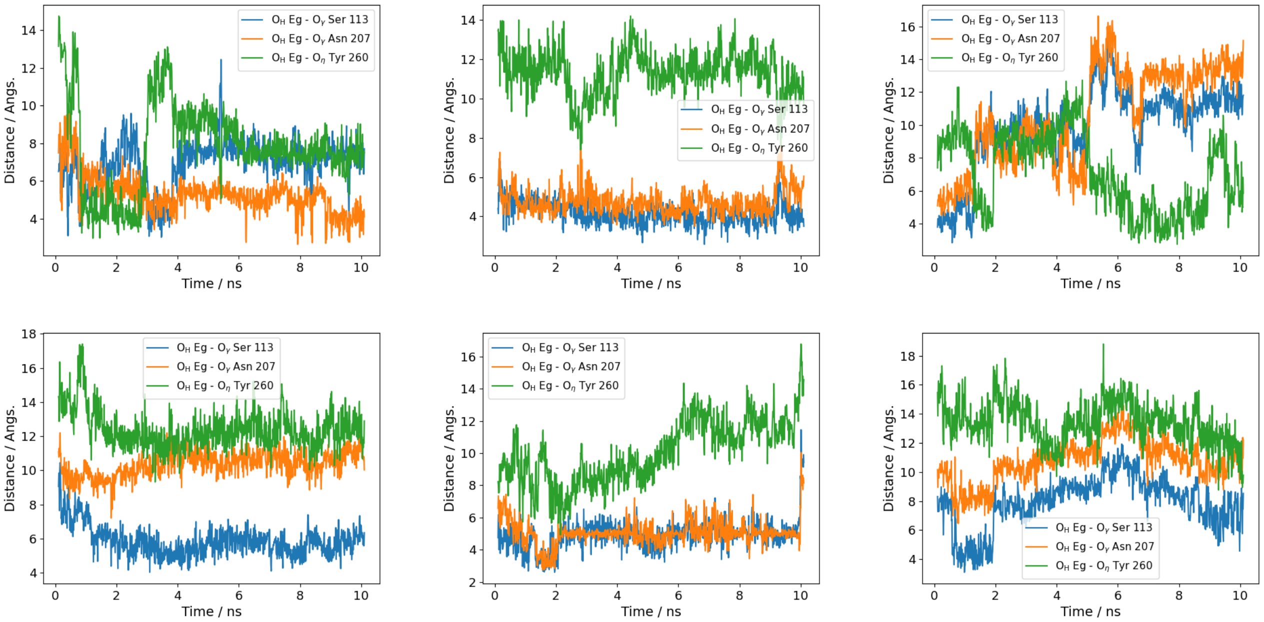


**Supplementary Figure S2** (conitued)

### Supplementary Figure S3

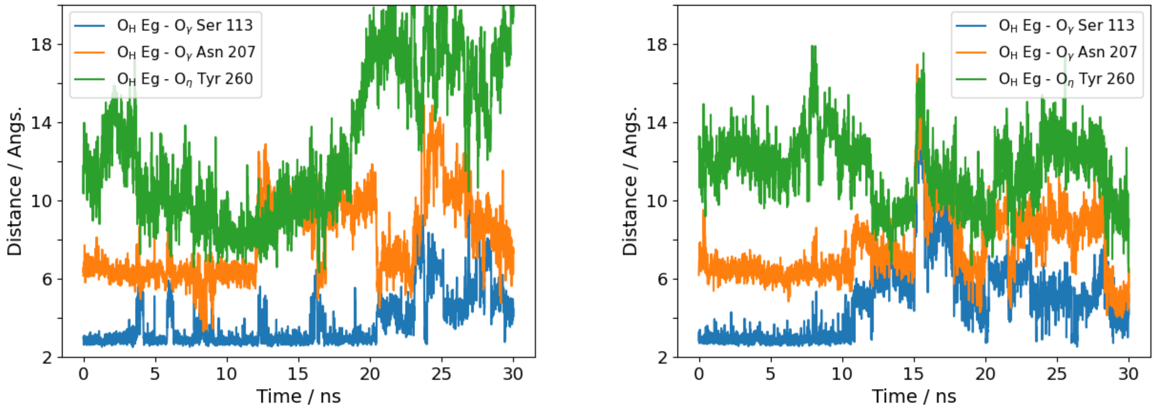


**Supplementary Figure S3** Two trajectories of the distance between $O_H$ atom of eugenol and O atoms of Ser113, Asn207, and Tyr260 that maintained the eugenol-Ser113 hydrogen-bond for longer than 10 ns.

### Supplementary Figure S4

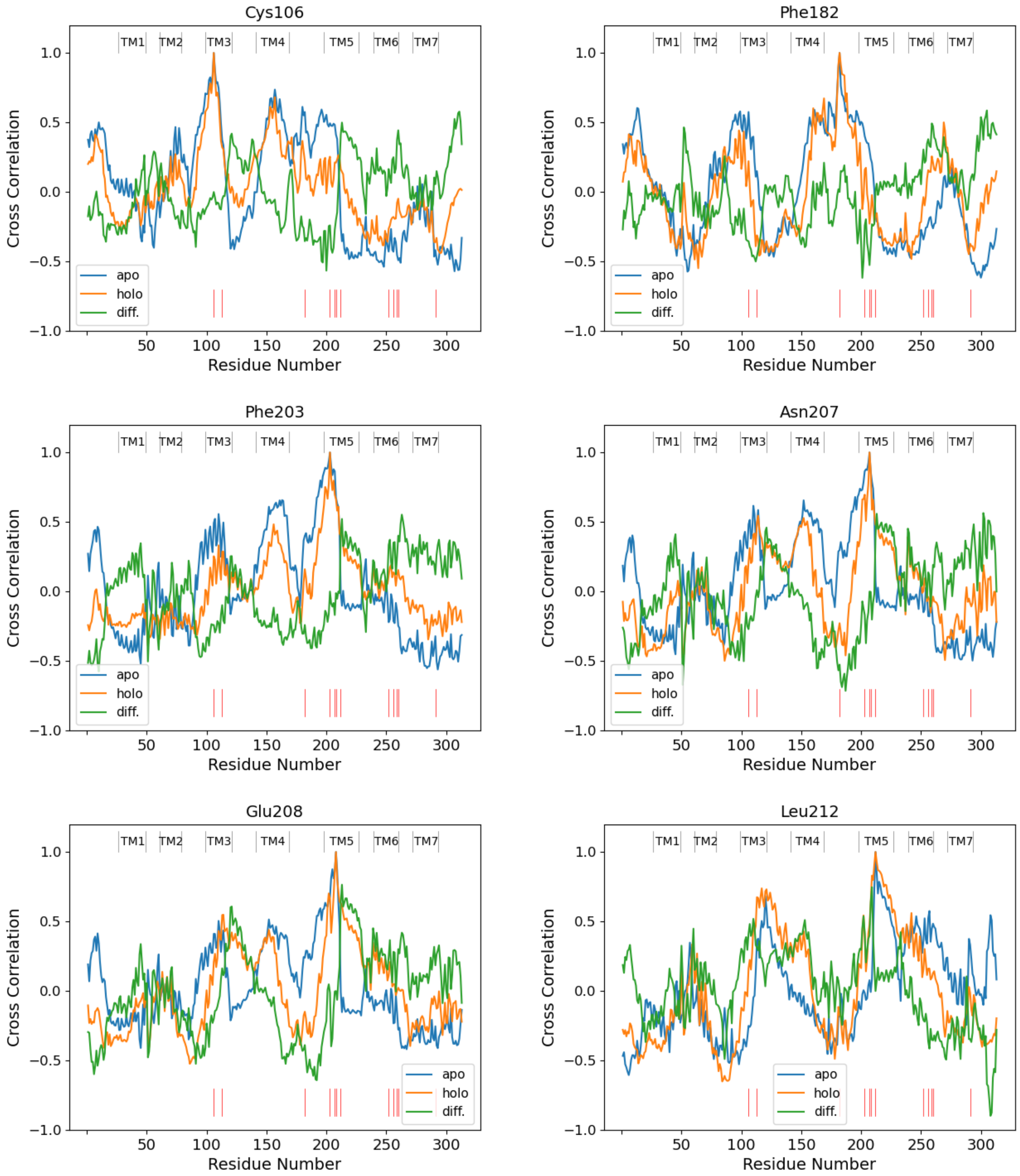


**Supplementary Figure S4** Sections of the dynamic cross-correlation maps (Figure 6 of the main text) at eleven key residues other than Ser113 displayed in Figure 7 (a) of the main text. (Continue to next page.)

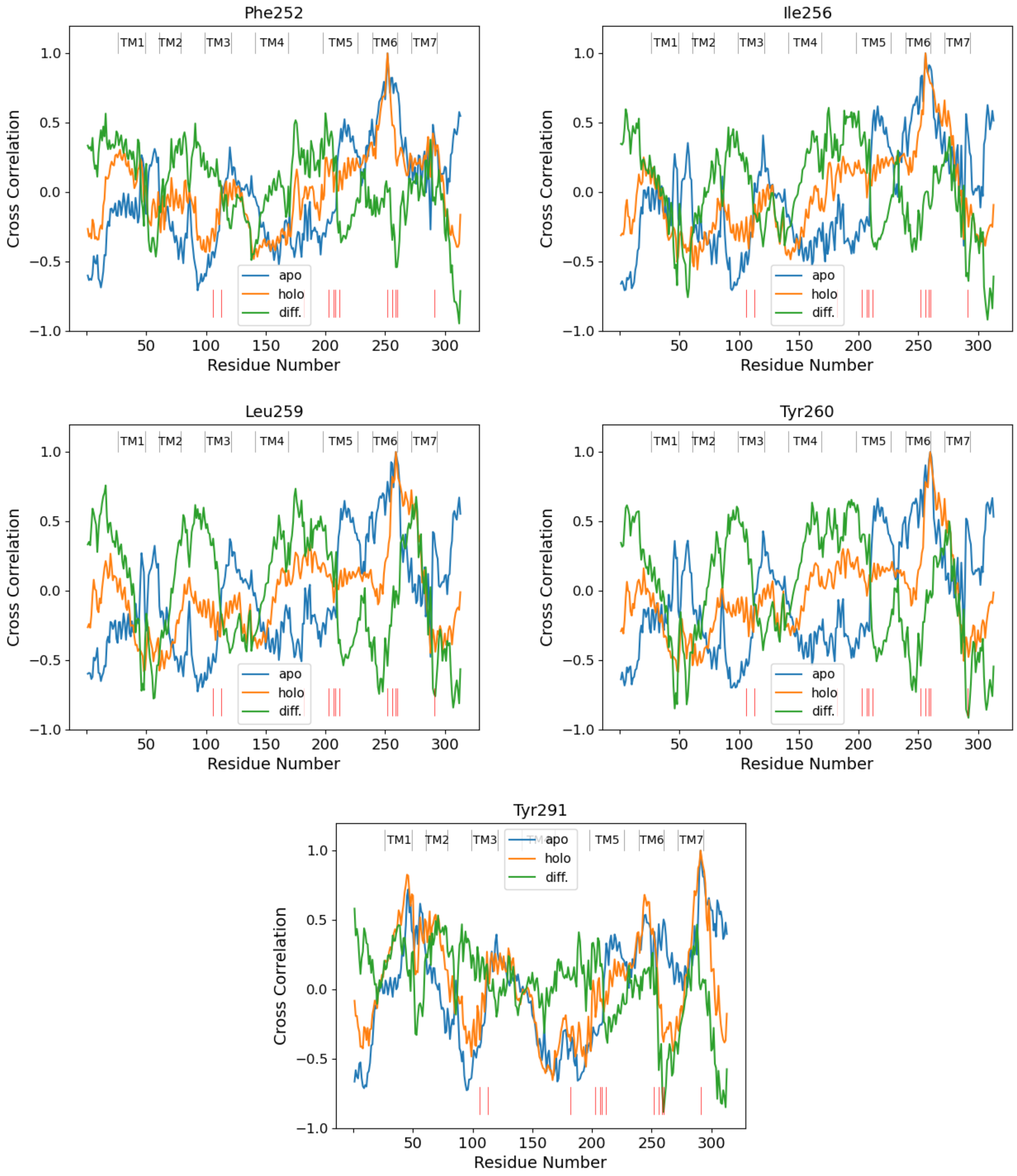


**Supplementary Figure S4** (continued)
